# Translational modulation by ISRIB alleviates synaptic and behavioral phenotypes in Fragile X syndrome

**DOI:** 10.1101/2023.09.08.553269

**Authors:** Rochelle L. Coulson, Valentina Frattini, Caitlin E. Moyer, Jennifer Hodges, Peter Walter, Philippe Mourrain, Yi Zuo, Gordon X. Wang

## Abstract

Fragile X syndrome (FXS) is caused by the functional loss of Fragile X messenger ribonucleoprotein (FMRP), a translational regulator that binds the transcripts of many proteins involved in synaptic function and plasticity. Dysregulated protein synthesis is a central effect of FMRP loss and to date direct translational modulation has not been leveraged in the treatment of FXS. Thus, we examined the effect of the small molecule translational modulator, Integrated Stress Response Inhibitor (ISRIB), in treating synaptic and behavioral symptoms of FXS. Using a combination of population-level single synapse analysis, *in vivo* two-photon microscopy, and behavioral analysis, we revealed a mechanism for the appearance of dense and immature dendritic spines, which are observed in human FXS patients and *Fmr1* knockout (KO) mice. Our data suggest that FMRP loss at the synapse dysregulates synaptic protein abundance and that spines are disproportionately stabilized through increased PSD-95 expression but prevented from maturing through reduced synaptic accumulation of glutamate receptors. ISRIB rescues these molecular and synaptic deficits in FXS and improves social recognition in *Fmr1* KO mice. These findings highlight the therapeutic potential of targeting core translational mechanisms and the differential impact of FMRP on synaptic and cellular function in FXS and neurodevelopmental disorders more broadly.

## Introduction

Neurodevelopmental disorders such as Autism spectrum disorder (ASD) and Fragile X syndrome (FXS) are associated with social and cognitive deficits, and are increasingly prevalent, with an estimated 1 in 54 children diagnosed with ASD in the United States (Maenner et al., 2020). FXS is the most common monogenic cause of inherited intellectual disability and ASD (Kelleher and Bear, 2008), and is characterized by many common autistic traits including cognitive dysfunction, social phobia, stereotyped behavior, hyperactivity, and hypersensitivity to sensory stimuli (Ey et al., 2011; Lozano et al., 2014). Due to the well-defined genetic etiology of FXS, pharmacological targeting of abnormal cellular and molecular pathways is possible and holds promise for the development of therapies for the treatment of neurodevelopmental disorders and ASD in general (Yamasue et al., 2019).

In humans, FXS is most often caused by the expansion of a CGG trinucleotide beyond 200 repeats within the 5’-untranslated region (UTR) of the Fragile X messenger ribonucleoprotein 1 gene (*FMR1)*. This inhibits *FMR1* transcription and the production of the highly conserved Fragile X messenger ribonucleoprotein (FMRP) (O’Donnell and Warren, 2002). FMRP is an RNA-binding protein that regulates the translation of approximately 4% of all transcripts in the brain (Brown et al., 2001). Many of these transcripts encode proteins that regulate synaptic function and plasticity, such as PSD-95, GluA1, GluA2, Arc, and Map1b (Antar et al., 2006; Bassell and Warren, 2008; Brown et al., 2001; Darnell et al., 2001; Dictenberg et al., 2008; Feng et al., 1997b, 1997a; Miyashiro et al., 2003; Weiler et al., 2004; Zalfa et al., 2007). Thus, loss of FMRP results in synaptic dysregulation, including protein synthesis-dependent processes such as group 1 mGluR-elicited long-term depression (mGluR-LTD) (Huber et al., 2002) and metabolic stress response (Didiot et al., 2009; Mazroui et al., 2002).

Protein translation, especially in the highly active awake brain, is extremely metabolically demanding. Behaviors such as socialization and learning induce significant metabolic load on a cellular and synaptic level in neurons (Attwell and Laughlin, 2001; Clarke and Sokoloff, 1999). Translation dependent mechanisms regulating this metabolic response are critical for proper synaptic network function and plasticity (Buffington et al., 2014).

Altered brain metabolism due to dysregulated protein synthesis in the absence of FMRP (Davidovic et al., 2011; El Bekay et al., 2007) suggests that drugs targeting translational regulation of stress pathways may alleviate FXS synaptic deficits and improve behavioral outcomes. One such drug is the small molecule Integrated Stress Response Inhibitor (ISRIB). ISRIB modulates translation in a context-specific manner, acting on a specific set of mRNAs that are preferentially translated under stress conditions, rather than altering bulk translation throughout the cell. By blocking this translational shift, ISRIB inhibits the translation-dependent phase of mGluR-LTD, which is also a key function of FMRP (Anand and Walter, 2020; Di Prisco et al., 2014; Hou et al., 2006; Koekkoek et al., 2005; Nakamoto et al., 2007; Nosyreva and Huber, 2006; Rabouw et al., 2019; Sidrauski et al., 2015; Wong et al., 2018). This suggests that local translational programs in neuronal synapses represent a key switching mechanism that regulates plasticity. In a normal neuron, FMRP regulates translation at the synapse (Weiler et al., 2004). In the absence of FMRP, ISRIB may compensate for some of the functions of FMRP as a translational regulator of plasticity and function at neuronal synapses. In support of this hypothesis, ISRIB prevents mGlu receptor agonists from eliciting mGluR-LTD and AMPA receptor internalization in cultured hippocampal neurons (Di Prisco et al., 2014). The efficacy of ISRIB in the absence of FMRP is unknown, thus we examined the effects of ISRIB treatment by characterizing the synaptic mechanisms underlying ISRIB activity in the *Fmr1* KO mouse model. Specifically, we sought to determine if ISRIB could compensate for the loss of translational regulation by FMRP and whether restoring synaptic protein expression could rescue behavioral outcomes in FXS.

In this study we examine the effects of FMRP deficiency at the molecular, synaptic, and behavioral levels to characterize the mechanisms underlying the cognitive impairment observed in FXS. We used synapse-scale proteomic imaging (Micheva and Smith, 2007; Wang et al., 2016, 2014) and *in vivo* dendritic spine imaging (Hodges et al., 2017) to identify previously undescribed molecular and structural synaptic changes caused by the loss of FMRP and their response to ISRIB. We further demonstrate that molecular synaptic rescue corresponds with a normalization of dendritic spine dynamics and social recognition. In summary, we show that the loss of FMRP, and the accompanying translational modulation at the synapse, induces a heterogenous synaptic response and dysregulated protein accumulation in synapses along with social deficits in *Fmr1* KO mice, which can be rescued through the restoration of synaptic protein accumulation by ISRIB.

## Results

### *Fmr1* KO mice exhibit increased PSD-95 accumulation in a subset of synapses

In humans and mouse models, FXS is associated with increased spine density and immature spine morphology (Comery et al., 1997; Irwin et al., 2000). These spine changes are hypothesized to be physical manifestations of synaptic network alterations (Grossman et al., 2006). However, the exact nature of synaptic protein changes in FXS has been difficult to assess, especially on a synapse-specific, population level. An important synaptic protein for studying the mechanism of excitatory synaptic network changes in FXS is PSD-95. PSD-95 plays a key role in organizing synaptic structure (Bats et al., 2007; Cane et al., 2014) and activity-dependent stabilization of spines and synaptic strengthening (De Roo et al., 2008). Further, PSD-95 mRNA stability and translation are regulated by FMRP (Muddashetty et al., 2011; Todd et al., 2003; Zalfa et al., 2007). We examined the effect of FMRP deficiency on PSD-95 abundance in individual synapses using a super-resolution, single synapse analysis method that combines array tomography (AT) with a computational synapse classification algorithm developed in our lab (Wang et al., 2016, 2014) to identify changes in specific molecularly-defined synaptic populations. This enables us to analyze all synapses of a specific class, e.g., vesicular glutamate transporter 1 (VGluT1)-expressing excitatory, cortical-cortical synapses, with single synapse resolution and metrics across entire populations (Wang et al., 2014). Thus, we were able to examine the unique effects of FMRP deficiency within specific synapse populations, while still maintaining our ability to resolve individual synaptic protein changes. We measured PSD-95 volume in cortical-cortical excitatory synapses. PSD-95 volume is demonstrated to be an excellent indicator of synapse size and maturity by electron microscopy (EM). In layer 1, the cumulative distribution plots of the wildtype (WT) and *Fmr1* KO PSD-95 volume size indicate a significant increase (p<2.2×10^−16^, Kolmogorov-Smirnov) in synapse size in layer 1 cortical-cortical synapses (Figure 1A). This shift is easily visualized in a density plot (Figure 1B), where the WT distribution of PSD-95 volume across the entire synapse population is more skewed towards smaller volumes than the *Fmr1* KO synapse population. This is further reflected in a 33% increase in median PSD-95 levels in vehicle-treated *Fmr1* KO mice compared to their vehicle-treated WT counterparts (p<2.2×10^−16^, Mann-Whitney U) (Figure 1A-C and Supplementary Data 1). In layer 2/3, the change in PSD-95 levels is minimal, with only a 3.8% increase in median volume observed in *Fmr1* KO mice (Supplementary Fig. 1 and Supplementary Data 1). Elevated PSD-95 levels in layer 1 *Fmr1* KO mouse synapses are reduced by ISRIB treatment, bringing them closer to WT levels (Figure 1A-C). Normalization of the distribution of PSD-95 across synapses suggests that the translational program disrupted by FMRP loss is rescuable by rebalancing translation with ISRIB.

**Figure 1.**
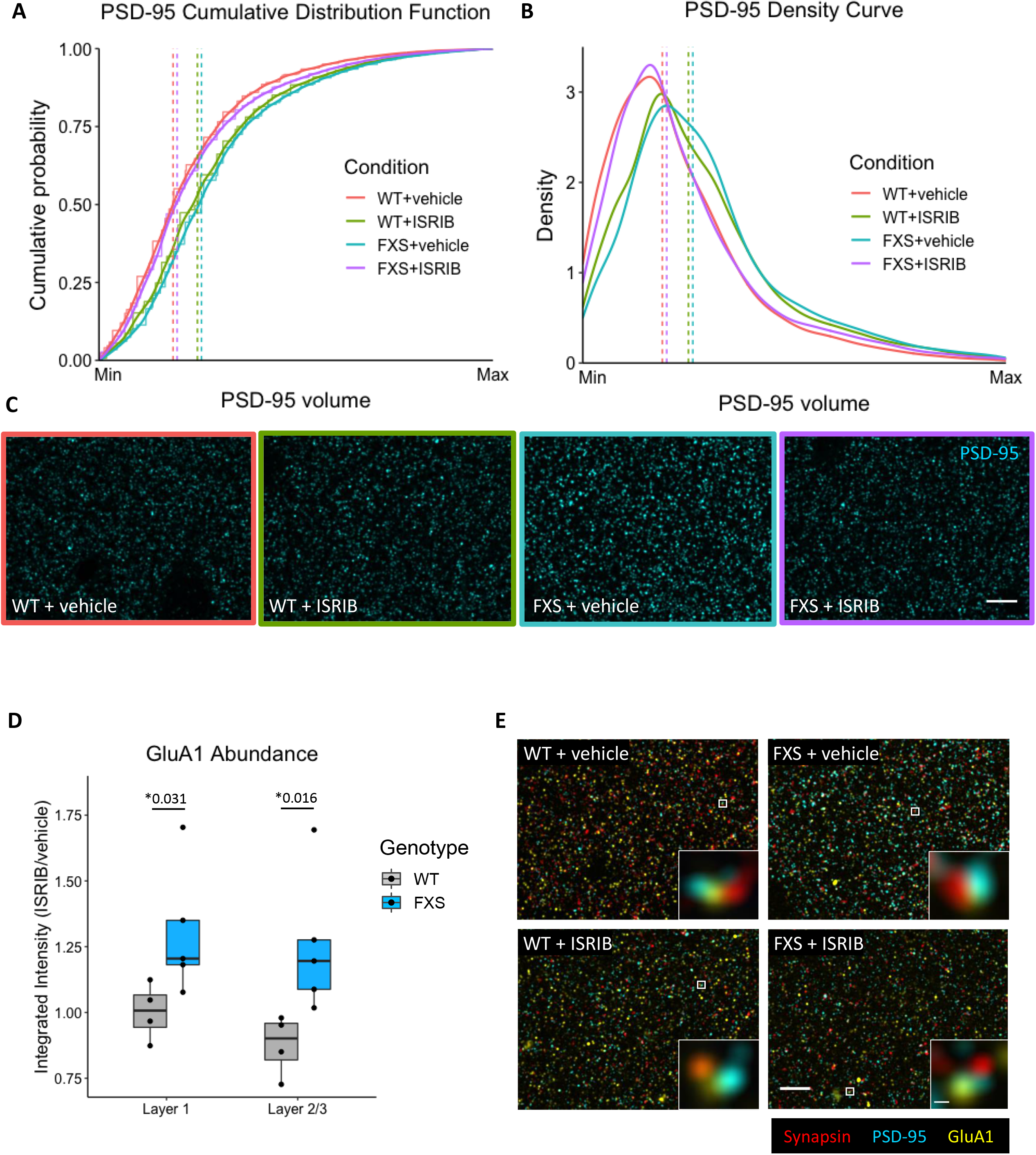
Array tomography (AT) analysis of PSD-95 and GluA1 abundance. (A) Cumulative distribution function (CDF) of PSD-95 size. PSD-95 volume was normalized to the maximum PSD-95 volume observed per mouse to normalize for experimental variability and the cumulative probability was plotted across the range of PSD-95 volumes. The raw CDF is shown as steps with the smoothed CDF overlayed for each condition. Vertical dashed lines indicate the median PSD-95 volume for each condition. Median = 0.21 WT+vehicle, 0.27 WT+ISRIB, 0.28 FXS+vehicle, 0.22 FXS+ISRIB. The distribution of PSD-95 among all synapses is altered in FXS layer 1 cortex, with fewer small PSD-95 synapses and a shift towards larger PSD-95 synapses. This shift results in a significantly altered PSD-95 synapse distribution (WT+vehicle vs FXS+vehicle, Kolmogorov-Smirnov, p<2.2×10^−16^) and a 33% increase in median PSD-95 size (WT+vehicle vs FXS+vehicle, Mann-Whitney U, p<2.2×10^−16^), which is decreased upon ISRIB treatment (FXS+vehicle vs FXS+ISRIB, Kolmogorov-Smirnov and Mann-Whitney U, p<2.2×10^−16^). (B) PSD-95 size shown as a density plot. Loss of small PSD-95 synapses is accompanied by a gain of larger PSD-95 synapses. Medians are shown as vertical dashed lines (values as stated in A). (C) Representative AT images for PSD-95 are shown for each condition. Imaging was performed blind to genotype and condition and images were taken at the same exposure. Images are a max projection throughout the depth of tissue. Scale bar = 5µm. FXS = *Fmr1* KO. n = 4 WT+vehicle (49,631 synapses), 4 WT+ISRIB (34,061 synapses), 5 FXS+vehicle (49,375 synapses), 5 FXS+ISRIB (49,350 synapses). (D) Integrated intensity for GluA1 was normalized for exposure and is the weighted average of VGluT1 and VGluT2 synapses. Tissues were processed and imaged in pairs and the ratio of integrated intensity for ISRIB/vehicle-treated mice was plotted. A ratio of one indicates no drug effect on abundance, with values above one indicating an increase in abundance with ISRIB treatment and values less than one indicating a decrease. ISRIB significantly increases GluA1 abundance in FXS synapses in both layers 1 (WT vs FXS, p=0.031) and 2/3 (WT vs FXS, p=0.016). (E) Representative AT images for layer 1 synapses in each condition including both a wide field view (scale bar = 5µm) and individual synapses (scale bar = 200nm). Individual synapses show Synapsin (red), PSD-95 (cyan), and postsynaptically localized GluA1 (yellow). Images are a max projection throughout the depth of tissue. Significance was calculated by Mann-Whitney U test. FXS = *Fmr1* KO. n = 4 WT+vehicle, 4 WT+ISRIB, 5 FXS+vehicle, 5 FXS+ISRIB.

### ISRIB increases GluA1 levels at the postsynaptic terminal in *Fmr1* KO mice

PSD-95 is important for the activity-dependent stabilization of spines and synaptic maturation (De Roo et al., 2008). The increase in volume of PSD-95 in layer 1 suggests a possible mechanism for the elevated stabilization of spines leading to higher spine and synapse density (Galvez and Greenough, 2005; Nimchinsky et al., 2001; Wang et al., 2014), but it does not provide a rationale for the immature morphology of the spines. A possible explanation is that loss of synaptic protein regulation in the absence of FMRP increases PSD-95 and spine stabilization but perturbs the accompanying process of spine and synapse maturation. One such maturation process proceeds through AMPA receptor subunit GluA1-mediated plasticity (De Roo et al., 2008; Ehrlich and Malinow, 2004; El-Husseini et al., 2000; Stein et al., 2003). PSD-95 accumulation is coupled with the clustering of GluA1 at the postsynaptic terminal (El-Husseini et al., 2000). The loss of FMRP-mediated translational regulation leads to reduction of membrane surface AMPA receptors localized at dendritic spines (Suresh and Dunaevsky, 2017). We hypothesized that translational normalization at the synapse by ISRIB could also normalize this glutamate receptor-mediated maturation. Therefore, we measured GluA1 protein levels at molecularly classified excitatory synapses in vehicle- and ISRIB-treated WT and *Fmr1* KO mice. To clearly determine the effect of ISRIB at the synapse, we calculated the ratio of GluA1 integrated intensity in excitatory synapses with or without ISRIB treatment (Figure 1D-E, Supplementary Fig. 2, and Supplementary Data 2). A ratio of one indicates no effect of treatment on GluA1 abundance, with values greater than one indicating an increase in protein abundance with ISRIB treatment, and values less than one representing a decrease in abundance. ISRIB treatment significantly increased postsynaptic GluA1 levels in *Fmr1* KO mice compared to WT in layers 1 and 2/3 of the cortex (Figure 1D) (p=0.03 (L1), p=0.02 (L2/3), Mann-Whitney U). We demonstrated that synaptic PSD-95 is elevated in *Fmr1* KO synapses which, without a concomitant increase in GluA1, may impede the normal process of synaptic strengthening and maturation. Furthermore, these FXS-specific synaptic protein changes can be normalized by ISRIB.

### ISRIB rescues abnormal spine dynamics in the cortex

The accumulation of PSD-95 and GluA1 in the postsynaptic density is involved in spine development and maturation (Bassell and Warren, 2008; El-Husseini et al., 2000). To further validate the synaptic specificity of ISRIB treatment in FXS, we next examined *Fmr1* KO spine dynamics and the effect of ISRIB in normalizing changes elicited by FMRP deficiency. Overabundance of dendritic spines is a neuropathological hallmark of FXS and is a well-established phenotype in mouse models of FXS (Bagni and Greenough, 2005; Comery et al., 1997; Hinton et al., 1991; Rudelli et al., 1985). *In vivo* imaging has revealed that excess dendritic spines found on layer 1 apical dendrites of layer 5 pyramidal neurons in the cortex of adult *Fmr1* KO mice are a consequence of an overproduction and stabilization of spines during adolescence (Hodges et al., 2017). To measure spine dynamics, we treated mice with ISRIB or vehicle control for four days and measured the rate of spine formation and elimination by transcranial two-photon microscopy in layer 1 of the cortex. Spine formation was significantly increased in *Fmr1* KO mice compared to WT mice with vehicle treatment (p=0.016, Mann-Whitney U). ISRIB treatment restored the rate of *Fmr1* KO spine formation to WT levels (WT + vehicle vs *Fmr1* KO + ISRIB, p=0.69, Mann-Whitney U). In contrast, the rate of spine elimination was not affected by FMRP deficiency and was unresponsive to ISRIB treatment (Figure 2 and Supplementary Data 3). This suggests an imbalance of spine stability in FXS that contributes to spine and synaptic changes in the disorder. Increased spine formation in FXS in conjunction with potential synaptic stabilization without a compensatory increase in spine elimination likely leads to the commonly observed phenotype of increased spine density in FXS.

**Figure 2.**
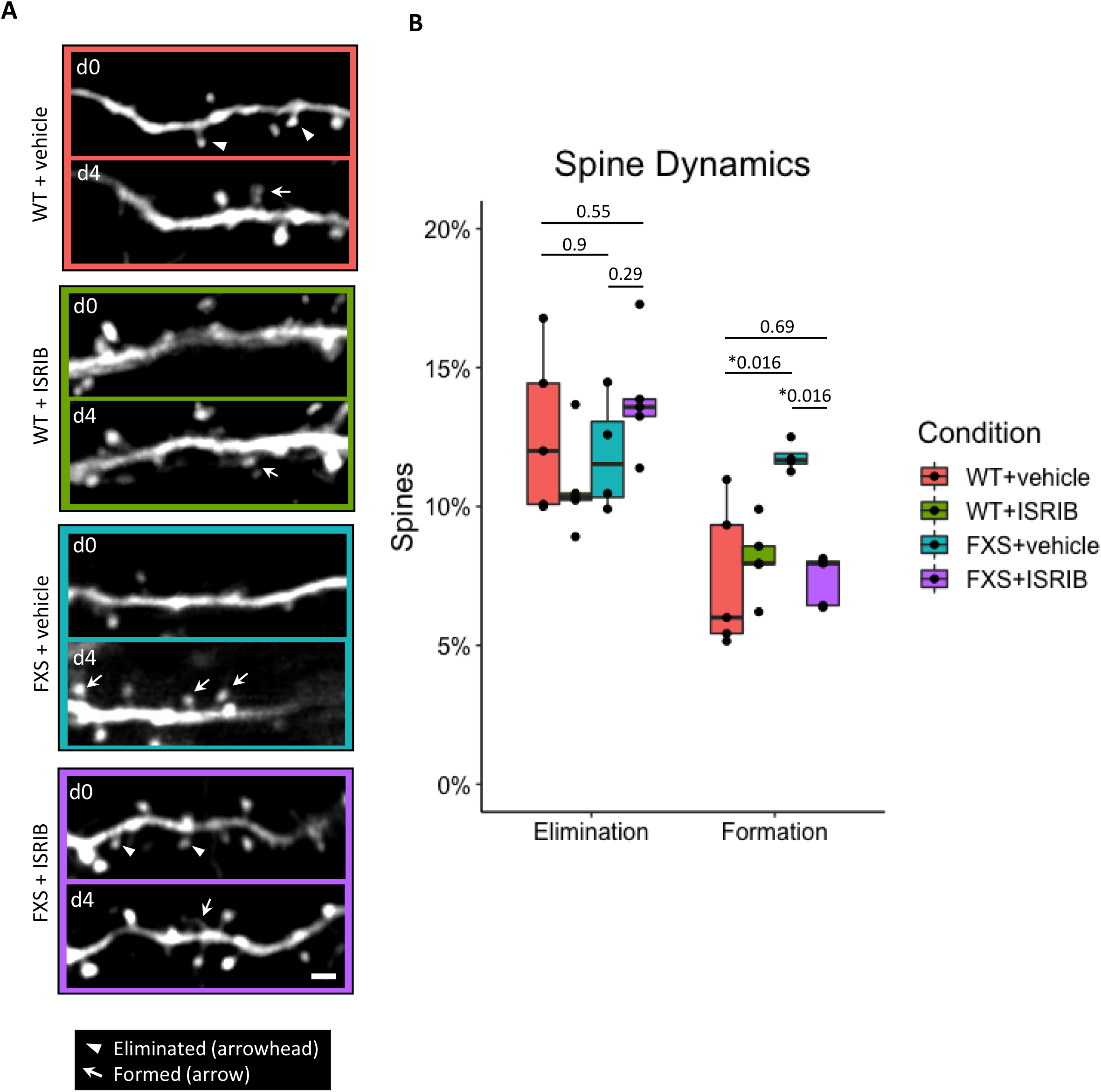
Transcranial two-photon imaging of dendritic spines in WT and *Fmr1* KO mice (FXS). (A) Representative images of dendritic spines at day 0 (d0) and day 4 (d4) for each condition. Spine formation is indicated by arrows and spine elimination is indicated by arrowheads. Scale bar = 2µm. Mice were treated with ISRIB or vehicle for four days and spines were imaged before (day 0) and after (day 4) treatment. Spine elimination and formation represents the change in spine count between day 0 and day 4. (B) Quantification of dendritic spine elimination and formation. Vehicle-treated FXS mice exhibit an aberrantly high rate of spine formation compared to vehicle-treated WT mice (WT+vehicle vs FXS+vehicle, p=0.016). ISRIB treatment restored the rate of FXS spine formation to WT levels (WT+vehicle vs FXS+ISRIB, p=0.69). No deficits were observed in spine elimination. Significance was calculated by Mann-Whitney U test. FXS = *Fmr1* KO. n = 5 WT+vehicle, 5 WT+ISRIB, 4 FXS+vehicle, 5 FXS+ISRIB.

### ISRIB ameliorates social recognition deficits in *Fmr1* KO mice

Similar to individuals with FXS, adolescent *Fmr1* KO mice exhibit impaired social interaction behavior (Kazdoba et al., 2014). Proper social interaction involves complex coordinated neural network function, and subtle deficits in cortical network function and connectivity can manifest as observable social interaction deficits. We thus sought to determine if the synaptic normalization we observed with ISRIB treatment is associated with improvements in behavioral outcomes. Using the three-chambered social novelty task, we asked whether ISRIB treatment improves social recognition in *Fmr1* KO mice (Figure 3A and Supplementary Data 4). *Fmr1* KO mice displayed a significant deficit in social novelty preference, spending an equal amount of time interacting with a familiar mouse and a novel mouse (WT + vehicle vs *Fmr1* KO + vehicle, p=0.011, Mann-Whitney U) (Dahlhaus and El-Husseini, 2010; McNaughton et al., 2008). ISRIB treatment increased social novelty preference in *Fmr1* KO mice, restoring preference to levels comparable to that of WT mice (*Fmr1* KO + vehicle vs *Fmr1* KO + ISRIB, p=0.019; WT + vehicle vs *Fmr1* KO + ISRIB, p=0.72, Mann-Whitney U). Conversely, WT and *Fmr1* KO vehicle and ISRIB-treated mice showed no difference in overall sociability (the preference to spend time with another mouse over an empty chamber) (Figure 3B and Supplementary Data 5). Behavioral effects were not attributable to changes in locomotor behavior, as there was no significant difference in the total distance traveled in the arena (Supplementary Fig. 3 and Supplementary Data 6). Overall, our results demonstrate that ISRIB treatment does rescue social recognition deficits in adolescent *Fmr1* KO mice.

**Figure 3.**
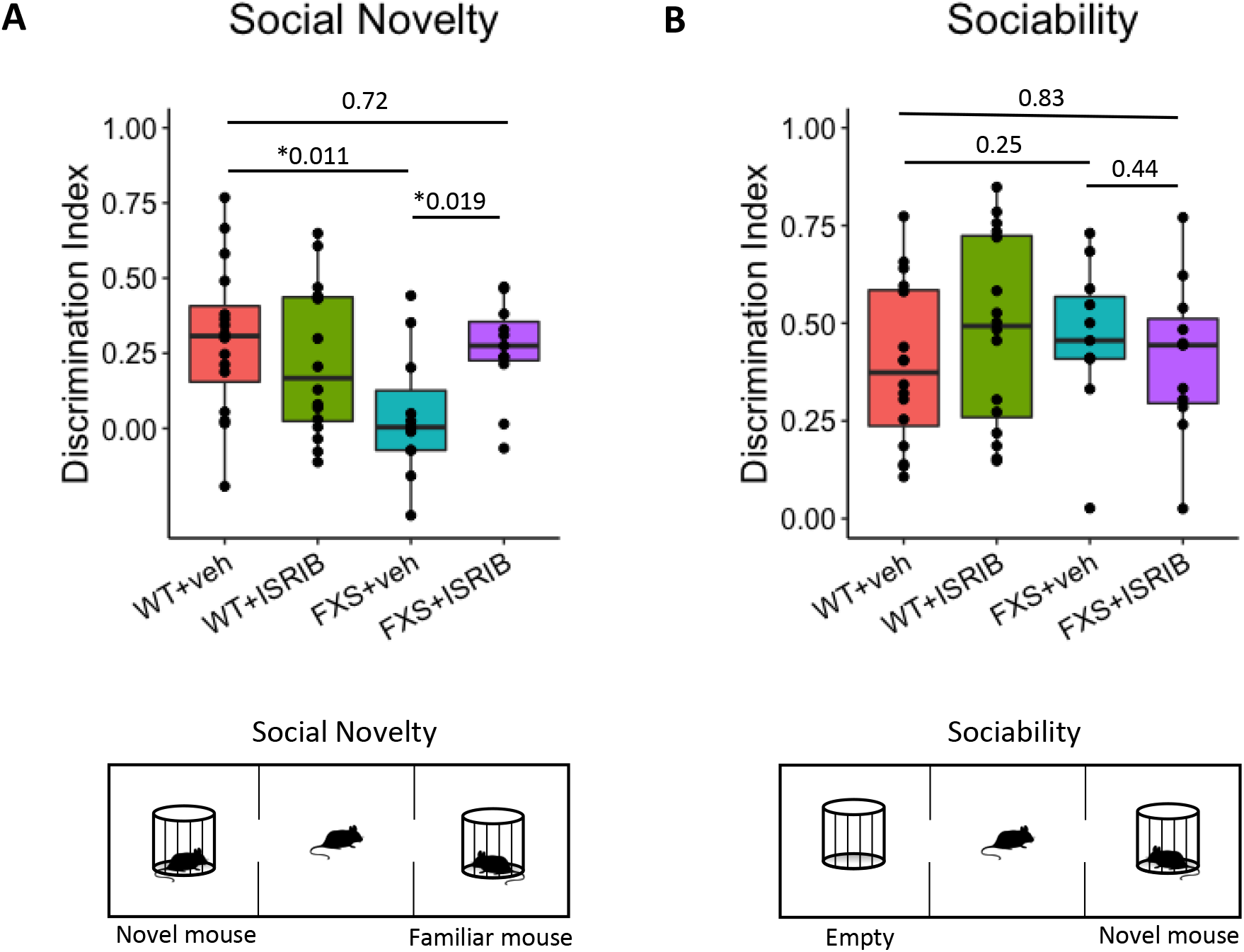
Behavioral analysis of *Fmr1* KO mice (FXS) with ISRIB treatment. (A) Social novelty was assessed using the three-chamber apparatus as diagrammed. One chamber houses a novel mouse, and the other chamber houses a familiar mouse. The amount of time spent investigating each mouse was quantified. The discrimination index is calculated as (novel interaction time -familiar interaction time)/(novel interaction time + familiar interaction time). FXS mice exhibit a deficit in social novelty preference compared to WT mice (WT+vehicle vs FXS+vehicle, p=0.011) and this effect is rescued by ISRIB treatment (FXS+vehicle vs FXS+ISRIB, p=0.019; WT+vehicle vs FXS+ISRIB, p=0.72). Significance was measured by Mann-Whitney U test. FXS = *Fmr1* KO. n = 16 WT+vehicle, 16 WT+ISRIB, 11 FXS+vehicle, 11 FXS+ISRIB. (B) Sociability was assessed using the three-chamber apparatus as diagrammed. One chamber houses an empty cage, and the other chamber houses a novel mouse. The amount of time spent investigating each cage was quantified and the discrimination index was calculated as described for social novelty. FXS mice showed no deficit in overall sociability (WT+vehicle vs FXS+vehicle, p=0.25). Significance was measured by Mann-Whitney U test. FXS = *Fmr1* KO. n = 16 WT+vehicle, 16 WT+ISRIB, 11 FXS+vehicle, 11 FXS+ISRIB.

## Discussion

In this study, we investigated a mechanism for the development of dense, immature spines in FXS and examined the therapeutic potential of translational reprogramming in FXS using the small molecule ISRIB. We demonstrated through population-level single synapse analysis that the loss of FMRP induces an unexpected shift towards increased synaptic PSD-95 and lower synaptic GluA1. This suggests a mechanism for the observation of increased density of spines with immature morphology in both individuals with FXS (Hinton et al., 1991; Rudelli et al., 1985) and *Fmr1* KO mice (Consortium, 1994). Our data suggest that loss of the translational regulator FMRP leads to increased spine stabilization through elevated levels of PSD-95 at the synapse, thus increased density of spines and synapses; while also preventing further accumulation of glutamate receptors, which results in immature synaptic and spine development (Figure 4)

**Figure 4.**
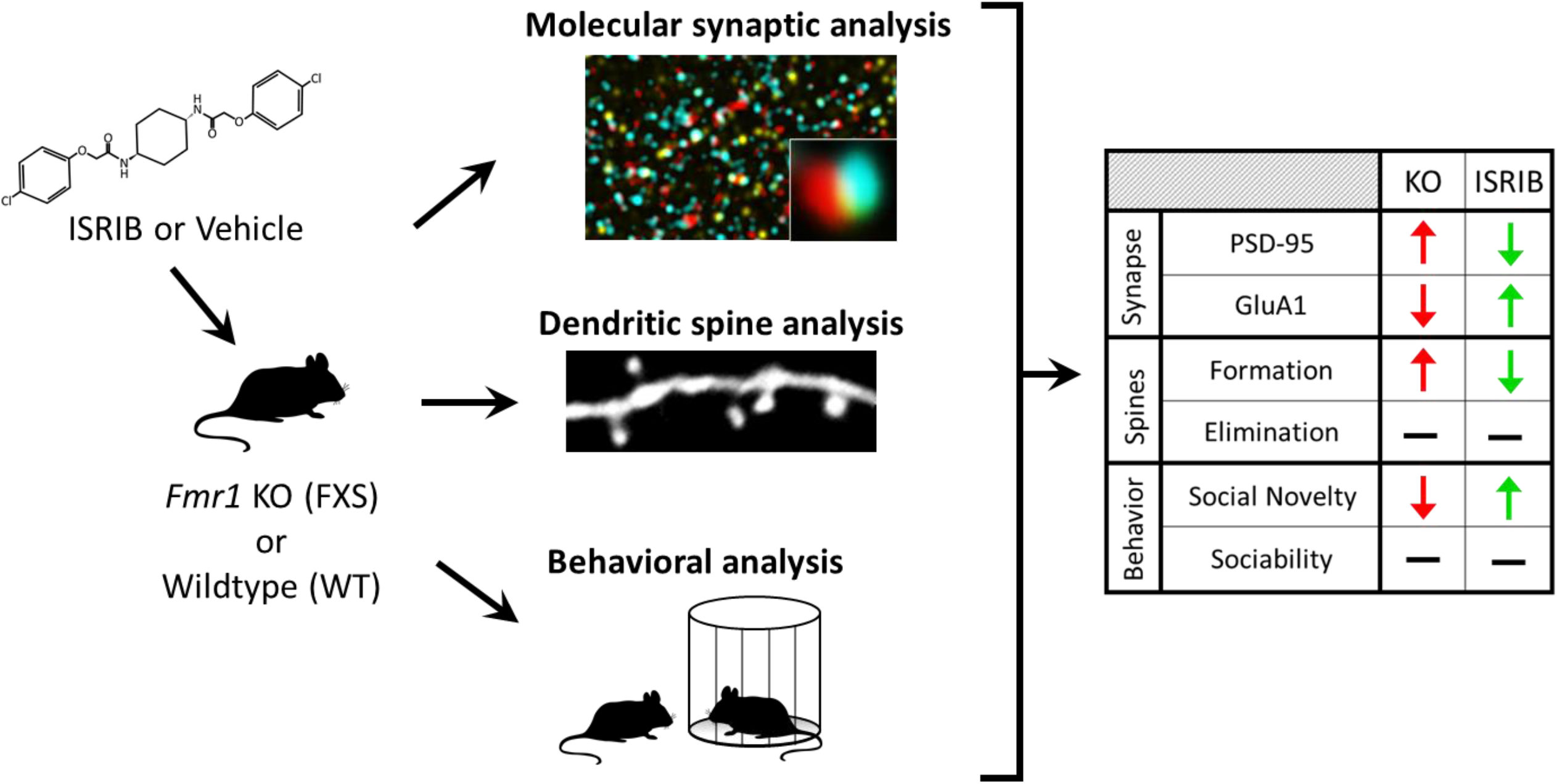
Summary of analyses and outcomes. *Fmr1* KO (FXS) and WT mice were treated with ISRIB or a vehicle control. The effects of FMRP loss and translational modulation by ISRIB treatment were assessed at the molecular, synaptic, and behavioral levels. FMRP loss at the synapse leads to a translational shift and the stabilization of aberrant immature spines with dysregulated PSD-95 and GluA1 levels. FXS mice also exhibit social recognition deficits characteristic of FXS. ISRIB restores synaptic translational control and rescues behavioral phenotypes.

Interestingly, the overabundance of spines in FXS is developmentally regulated, observed early during postnatal week 1, normalized in adolescence from weeks 2-4, and re-emerging in adulthood (Galvez and Greenough, 2005; He and Portera-Cailliau, 2013; Nimchinsky et al., 2001; Wang et al., 2014). Our study examined the developmental window in which spine abundance is normalized, thus we observed no defect in synaptic density in P26-P32 *Fmr1* KO cortex (Supplementary Fig. 4 and Supplementary Data 7). This apparent normalization belies developmental dynamics that will drive the nervous system towards aberrant connectivity. Our dendritic spine dynamics data is a window into this underlying developmental context. Due to FMRP loss-mediated decoupling of PSD-95 and GluA1 synaptic accumulation, dendritic spines exhibit increased stabilization leading to a large population of stabilized spines without a commensurate increase in elimination. This is likely the cause of the re-emergence of elevated immature spine density and synapses in the mature FXS brain. Thus, modulation of excessive spine formation with ISRIB treatment during this developmental window may mitigate the resulting overabundance of spines that emerges in adult *Fmr1* KO mice and improve cognitive and behavioral outcomes.

FMRP is a synaptic regulator of translation. Its function is controlled by synaptic activity, and it acts to regulate the translation of proteins involved in the development and maturation of spines and synapses (Antar et al., 2006; Bassell and Warren, 2008; Brown et al., 2001; Buffington et al., 2014; Darnell et al., 2001; Kelleher and Bear, 2008; Miyashiro et al., 2003; Zalfa et al., 2007). FMRP regulates translational programs at each synapse to tune activity-dependent processes in neural plasticity (Weiler et al., 2004). This allows a neuron to modulate and prioritize its incoming information, thus effectively increasing the operational dynamic range of the nervous system. The loss of FMRP critically reduces the ability of synapses to plastically respond to intercellular communication. A molecule that can restore, even partially, this synaptic control of translation could in theory compensate for some of the functions of FMRP and return the synaptic network to near normal function. We proposed that ISRIB, a stress-mediated translational modulator, might function as a compensatory mechanism in the absence of FMRP at the synapse.

ISRIB belongs to a class of small molecules that can modulate translation at the initiation phase, much like FMRP, and reduce the translation of specific classes of mRNAs (Sidrauski et al., 2013; Tsai et al., 2018). Our study showed that ISRIB indeed can improve molecular accumulation of synaptic proteins, dendritic spine dynamics, and behavioral function in a mouse model of FXS (Figure 4). ISRIB improves social recognition memory, rescuing social novelty preference in *Fmr1* KO mice. Behavioral outcomes are a key functional measure of underlying improvements in molecular and cellular function, which often impact central regulatory pathways. Previously, ISRIB has been shown to improve cognitive function in brain injury (Chang et al., 2021; Chou et al., 2017), cancer (Lorenz et al., 2021), neuropsychiatric disorders (Kabir et al., 2017), neurodegenerative disorders (Oliveira et al., 2021; Wong et al., 2018; Xu et al., 2021), and even healthy cognition (Sidrauski et al., 2013). Our study demonstrated the beneficial effects of ISRIB in a mouse model of FXS, which in combination with recent studies showing the impact of ISRIB treatment in models of Down syndrome (Zhu et al., 2019) and MEHMO (Mental retardation, Epileptic seizures, Hypogonadism and Hypogenitalism, Microcephaly, and Obesity), a rare X-linked disorder (Young-Baird et al., 2020), expands its therapeutic potential to neurodevelopmental disorders. This broad efficacy of ISRIB treatment suggests a potentially shared framework of dysregulated translation across neurological diseases. Most therapeutic approaches to FXS target downstream of the core translational impact of FMRP loss. We demonstrate that by targeting one of the direct effects of FMRP loss, synaptic translational dysregulation, we can rescue synaptic and behavioral phenotypes. These observations suggest that targeting this pathway with ISRIB may represent a therapeutic entry point for addressing the pathophysiology underlying FXS, with potential for alleviating the impact of FMRP deficiency across the brain.

## Acknowledgements

This work was supported by the National Institutes of Health F32 HD103451 (RLC), F32 ES026872 (CEM), K01 AG061230 (GXW), R01 NS104950 (PM and YZ), R01 MH109475 (YZ), R01 AG071787 (YZ), FRAXA Research Foundation (RLC, GXW, and PM), John Merck Foundation (GXW and PM), Brain and Behavior Research Foundation (GXW), and Stanford (RLC and GXW). PW is an HHMI investigator.

We thank Sabyn Nopar and Keerat Bains for help with animal husbandry, mouse perfusions, and necroscopies and Ana Radonovich for help with 3-chamber tests.

CEM contributed to the project consistent with her previous role as a postdoctoral scholar at UCSC. The views and opinions expressed in this manuscript are those of the authors only and do not necessarily represent the views, official policy or position of the U.S. Department of Health and Human Services or any of its affiliated institutions or agencies.

## Author contributions

RLC, CEM, JH, PW, PM, YZ, and GXW conceived and designed the project; RLC, VF, CEM, and JH performed experiments and analyzed data; RLC, CEM, and GXW wrote the paper; all authors edited and approved the paper.

## Methods

### Experimental Animals

The Institutional Animal Care and Use Committee of University of California Santa Cruz approved all animal care and experimental procedures. Mice were group housed under a 12-hour light-dark cycle with *ad libitum* food and water. The *Fmr1* KO mice were a gift from Dr. Stephen T. Warren at Emory University. Thy1-YFP-H transgenic mice (stock number 003782), used for dendritic spine experiments were purchased from The Jackson Laboratory (Bar Harbor, ME). All mice were backcrossed with C57BL/6 mice more than 10 generations to produce congenic strains. Males (*Fmr1* KO and WT littermates) were used in all experiments. Experiments were conducted on adolescent mice, age P26-P32 for all experiments.

### Drug treatment

Animals received ISRIB at 2.5 mg/kg in 50% DMSO and 50% PEG400 for two-photon microscopy or 1 mg/kg in 5% DMSO, 2% Polysorbate 80, 20% PEG400, and 73% dextrose water for array tomography and behavioral testing, or a matched volume of the appropriate vehicle once per day by intraperitoneal injection. Mice were treated for 4 days for array tomography and two-photon imaging and 2 days for behavioral testing.

### Array tomography

Tissue and ribbon array preparation, immunohistochemistry, and microscopy were performed as described previously (Micheva and Smith, 2007), and outlined below.

#### Tissue and ribbon array preparation

Tissue was microwave fixed in 4% paraformaldehyde. A small piece of tissue (∼2mm deep x 1mm wide x 1mm long) was cut from the motor cortex of WT and *Fmr1* KO males age P26-P32. Tissue was dehydrated in 50%, 70%, 95% ethanol, then embedded in LRWhite resin (hard grade) in gelatin capsules. Sections of 70nm were cut using an ultramicrotome, creating a ribbon array of serial sections, which was placed on a gelatin-coated coverslip. Tissues were paired so that each coverslip contained two ribbons, one of each genotype for a given treatment (vehicle or ISRIB), to minimize slide to slide variability.

#### Immunohistochemistry

For immunostaining, coverslips containing ribbon arrays were washed in Tris buffer (Tris/50mM glycine/0.05% Tween) and primary antibodies were applied in Tris buffer containing 0.1% BSA. Coverslips containing antibodies were incubated at 4°C overnight in a humidified chamber. Coverslips were washed in Tris buffer 3×5min and secondary antibodies were applied in Tris buffer containing 0.1% BSA for 30min at room temperature. Coverslips were washed with Tris buffer 3×5min, rinsed with milliQ H2O, and mounted in SlowFade Gold anti-fade with DAPI (Invitrogen). After imaging, mounting media was washed with milliQ H2O and antibodies were eluted with 0.2M NaOH/0.02% SDS for 5min. The next round of staining was performed immediately after elution or coverslip was allowed to dry completely.

| **Antibody** | **Source** | **Identifier** |
| --- | --- | --- |
| GluA1 | Millipore | Cat# AB1504; RRID: AB_2113602 |
| VGluT2 | Millipore | Cat# AB2251; RRID: AB_1587626 |
| PSD-95 | Cell Signaling | Cat# 3450; RRID: AB_2292883 |
| VGluT1 | Millipore | Cat# AB5905; RRID: AB_2301751 |
| Synapsin-1 | Cell Signaling | Cat# 5297; RRID: AB_2616578 |
| Goat anti-Mouse Alexa 594 | Invitrogen | Cat# A110032 |
| Goat anti-Guinea Pig Alexa 594 | Invitrogen | Cat# A11076 |
| Goat anti-Mouse Alexa 488 | Invitrogen | Cat# A11029 |
| Goat anti-Rabbit Alexa 488 | Invitrogen | Cat# A11034 |
| Goat anti-Rabbit Alexa 647 | Invitrogen | Cat# A21245 |
| Goat anti-Mouse Alexa 647 | Invitrogen | Cat# A21236 |

#### Microscopy

Ribbons were imaged on a Zeiss Axio Imager.Z1 Upright Fluorescence Microscope with a motorized stage and Axiocam HR Digital Camera. The same field of view for each Layer 1 and Layer 2/3 cortical region were imaged for every section within each array and for every round of staining. Images were taken using a Zeiss 63x/1.4 NA Plan Apochromat objective with the same exposure for both ribbons on each coverslip. All microscopy was performed blinded to genotype and treatment.

#### Image processing

Image stacks were imported into FIJI and registered across all image sessions using a rigid model with interpolation. Registered images were deconvolved in Matlab using the Richardson-Lucy technique with background subtraction and an empirical point spread function (PSF) through ten iterations. Empirical PSF were measured using the actual imaging system with 110nm beads mounted on slides. Images were aligned with TrakEM in FIJI by both rigid least squares (linear feature correspondences) and elastic (non-linear block correspondences) alignments.

#### Synapse classification

Synapse classification is based on a set of requirements, as described previously (Allen et al., 2012; Hiu et al., 2016; Wang et al., 2014). Utilizing specific protein localization profiles for each synaptic population, established by Wang et al., 2016, synapses are identified and classified using three points: the origin (PSD-95), the terminus (Synapsin), and a third marker with a defined pre-or post-synaptic distance (VGluT1 or VGluT2) (Wang et al., 2016). A vector is drawn between the centers of mass from origin to terminus, defining a colocaliztion sphere centered around the origin. A third marker, such as VGluT1 or VGluT2 must fall within this sphere. Distances to the pre- and post-synaptic marker are utilized to confirm potential synapses identified by this PSD-95/Synapsin colocalization and provide more specificity in synapse type. For a presynaptic marker, the synapse is verified if the distance to Synapsin is smaller than the distance to PSD-95, otherwise, the potential synaptic axis is removed. If more than one third point falls within a colocalization sphere, the dataset is checked for overlapping spheres and points are assigned to maximize verified synapses. If a single third point is shared between two spheres, it is assigned to the synapse with the shortest pre-or post-synaptic distance as previously defined. Synaptic density is calculated as the total number of cortical-cortical excitatory synapses per non-nuclear volume of tissue.

#### Synaptic protein analysis

Signal volume represents the number of voxels per puncta. Volume was used as a measure of PSD-95 abundance and postsynaptic size for each individual synapse, and the distribution was plotted as a cumulative distribution function and density curve.

Significant shifts in distribution were tested by Kolmogorov-Smirnov and significant differences in median volume were tested by Mann-Whitney U. Integrated intensity was used as a measure of GluA1 abundance. GluA1 abundance is a weighted average of VGluT1 and VGluT2 synapses based on their relative contribution to the excitatory synapse population (L1: 89.144% VGluT1, 10.856% VGluT2; L2/3: 85.008% VGluT1, 14.992% VGluT2). The ratio of ISRIB/vehicle was calculated for each pair of tissue imaged at the same time for each genotype. Significant differences were tested by Mann-Whitney U.

#### Normalization

All tissues were processed in pairs (fix, dissect, embed, section, and stain). Exposure levels and segmentation thresholds were set with identical parameters based on previously validated methods. The pairing of animals and tissues allows us to reduce variability and minimize animal use. For ratio analyses (eg. ISRIB/vehicle), pair information is maintained, and direct comparisons are made. When pairs are mixed (eg. distribution curves), normalization of the entire distribution is performed on a per animal basis based on each animal’s maximum observed value, correcting for shifts in distribution bounds.

### In vivo dendritic spine imaging

Transcranial two-photon imaging of dendritic spines and data analysis were conducted as described previously (Hodges et al., 2017). Tg(Thy1-YFP)H were crossed into the *Fmr1* KO line to produce a line expressing YFP within the apical dendrites and spines of motor cortex layer 5 neurons, making them amenable to transcranial two-photon imaging of spine dynamics. P26-P32 YFP-expressing WT and *Fmr1* KO males were used for *in vivo* two-photon imaging experiments. Collected image stacks were analyzed using ImageJ. Mice received daily vehicle or ISRIB treatment for 4 days between the day 0 and day 4 imaging sessions. The percentage of eliminated and formed spines was calculated as the difference in the number of spines between the day 0 and day 4 imaging sessions divided by the number of spines counted for the day 0 imaging session. Mann-Whitney U test was used to determine differences in spine dynamics.

### Three-chamber tests

The same general testing protocol was used to evaluate both sociability and social novelty in vehicle and ISRIB-treated P26-P32 *Fmr1* KO mice and WT littermates. The test apparatus consists of three chambers (20 × 40 × 20 cm each) with openings to enable free access to each chamber (Kaidanovich-Beilin et al., 2011). The two side chambers contain cylindrical wire cages as confinements for novel and familiar age-matched, same sex probe mice. First, the mice were allowed to freely explore all three chambers for 10 minutes without any additional mice present in the wire enclosures.

During the sociability test phase, an unfamiliar C57Bl/6J mouse was put inside one wire cage while the other wire cage remained empty. During the social novelty test phase, an unfamiliar male C57Bl/6J mouse was placed in one wire cage, while the other wire cage contained a familiar mouse. During both testing phases, the subject mice were allowed to freely explore all chambers for 10 minutes. Chambers and cages were cleaned with 70% ethanol between each testing session. Mice were video recorded and the duration of time each mouse spent investigating each wire cage was quantified using Behavioral Observation Research Interactive Software (BORIS v. 7.10.2) (Friard and Gamba, 2016). Interaction is determined when the mouse is close to and facing the wired cage. Behaviors were annotated manually, and analysis was performed blind to genotype. Assessment of general locomotion was carried out by measuring the total distance traveled by each mouse during habituation and testing. Changes in social novelty, sociability, and locomotion were tested by Mann-Whitney U test.

## Supplemental Information

<u>Supplementary Figure 1</u>

AT analysis of synapse-level PSD-95 volume in layer 2/3 of the cortex. (A) Cumulative distribution function (CDF) of PSD-95 size. PSD-95 volume was normalized to the maximum PSD-95 volume observed per mouse to normalize for experimental variability and the cumulative probability was plotted across the range of PSD-95 volumes. The raw CDF is shown as steps with the smoothed CDF overlayed for each condition. Vertical dashed lines indicate the median PSD-95 size for each condition. Median = 0.22 WT+vehicle, 0.29 WT+ISRIB, 0.23 FXS+vehicle, 0.27 FXS+ISRIB. (B) PSD-95 size shown as a density plot. Medians are shown as vertical dashed lines (values as stated in A). FXS = *Fmr1* KO. n = 4 WT+vehicle (36,371 synapses), 4 WT+ISRIB (27,532 synapses), 5 FXS+vehicle (45,964 synapses), 5 FXS+ISRIB (48,654 synapses).

<u>Supplementary Figure 2</u>

Individual channels for representative wide field view AT images for layer 1 synapses shown in Figures 1C and 1E. Images are a max projection throughout the depth of tissue. Scale bar = 5µm. Synapsin = red, PSD-95 = cyan, GluA1 = yellow, VGluT1 = green, VGluT2 = magenta.

<u>Supplementary Figure 3</u>

Analysis of locomotor behavior during behavioral testing. Assessment of general locomotion was carried out by measuring the total distance traveled by each mouse over the course of the 10-minute habituation period or social novelty and sociability tests. WT and *Fmr1* KO mice show no difference in total distance traveled in the arena during behavioral testing. Significance was calculated by Mann-Whitney U test. FXS = *Fmr1* KO. Habituation n = 13 WT+vehicle, 15 WT+ISRIB, 9 FXS+vehicle, 10 FXS+ISRIB; Social Novelty n = 10 WT+vehicle, 14 WT+ISRIB, 5 FXS+vehicle, 8 FXS+ISRIB; Sociability n = 13 WT+vehicle, 15 WT+ISRIB, 9 FXS+vehicle, 10 FXS+ISRIB.

<u>Supplementary Figure 4</u>

AT analysis of synaptic density in layers 1 and 2/3 of the cortex for each condition. Synapses are identified and classified using three points: the origin (eg. PSD-95), the terminus (eg. Synapsin), and a third marker with a defined pre-or post-synaptic distance (eg. VGluT1, VGluT2). Synaptic density is calculated for the total non-nuclear volume. *Fmr1* KO mice do not exhibit altered synaptic density at P26-P32. Significance was calculated by Mann-Whitney U test. FXS = *Fmr1* KO. n = 4 WT+vehicle, 4 WT+ISRIB, 5 FXS+vehicle, 5 FXS+ISRIB.

<u>Supplementary Table 1</u>

PSD-95 volume per synapse. Raw volume measurements (voxels per puncta) are shown as well as scaled volume as a proportion of the maximum volume observed per mouse.

<u>Supplementary Table 2</u>

GluA1 normalized integrated intensity measurements. Measurements are a weighted average of the integrated intensity of VGluT1 and VGluT2 synapses based on their relative contribution to the excitatory synapse population (L1: 89.144% VGluT1, 10.856% VGluT2; L2/3: 85.008% VGluT1, 14.992% VGluT2).

<u>Supplementary Table 3</u>

Dendritic spine counts. Counts for WT and *Fmr1* KO mice with vehicle and ISRIB treatment at the beginning of treatment (day 0) are given as well as counts for spines that are stable, eliminated, or newly formed at the end of treatment (day 4).

<u>Supplementary Table 4</u>

Social novelty preference. Time spent with each mouse is shown in both seconds and as a percentage of total interaction time. Discrimination index (DI) is the difference between time spent with the novel and familiar mouse divided by the total time spent interacting with either.

<u>Supplementary Table 5</u>

Sociability. Time spent with either the empty cage or a mouse is shown in both seconds and as a percentage of total interaction time. Discrimination index (DI) is the difference between time spent with the mouse and empty cage divided by the total time spent interacting with either.

<u>Supplementary Table 6</u>

Locomotion during behavioral testing. The total distance traveled (cm) is recorded for each mouse during habituation, social novelty testing, and sociability testing.

<u>Supplementary Table 7</u>

Synaptic density. Density is calculated as the total number of cortical-cortical excitatory synapses per non-nuclear volume of tissue. This density is then reported per cubic micron.

