## Supplementary figures and images for "Translational modulation by ISRIB alleviates synaptic and behavioral phenotypes in Fragile X syndrome"

### Supplemental figures

Supplementary Figure 1

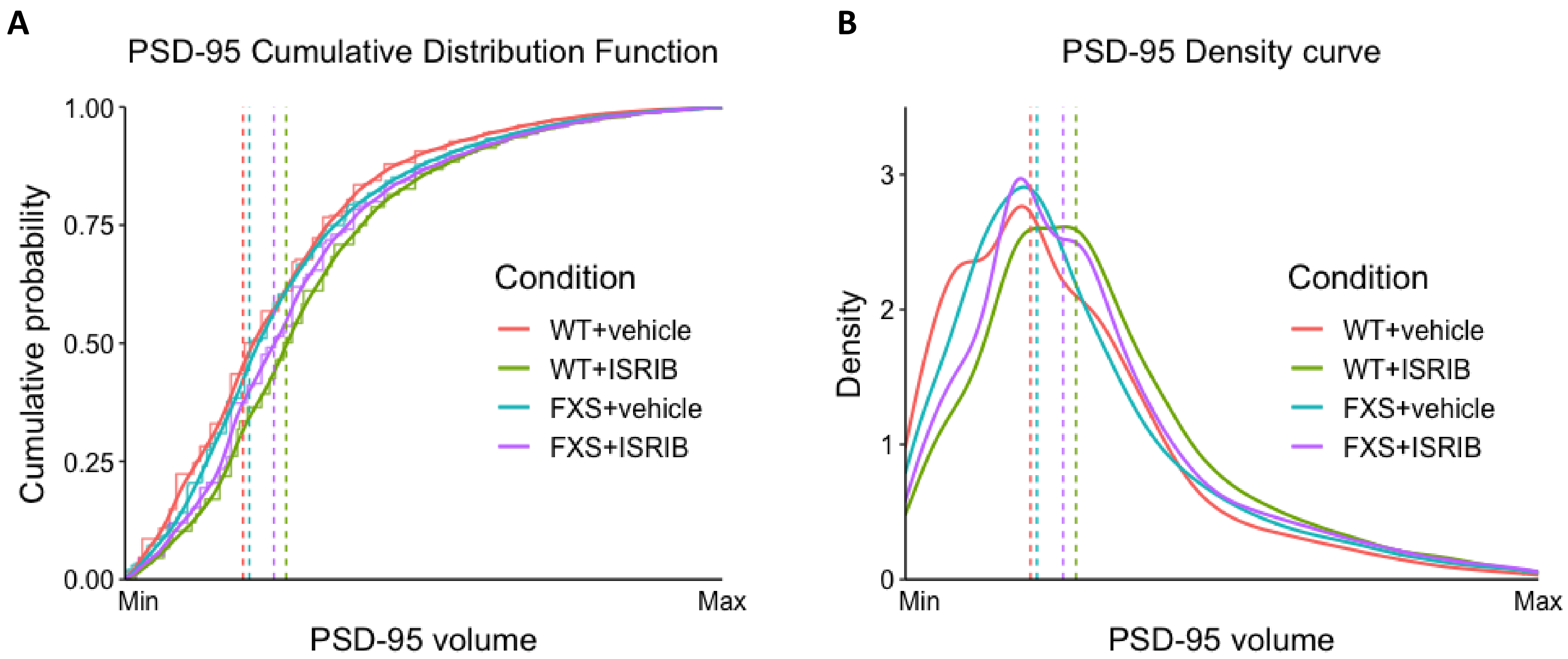

#### Supplementary Figure 2

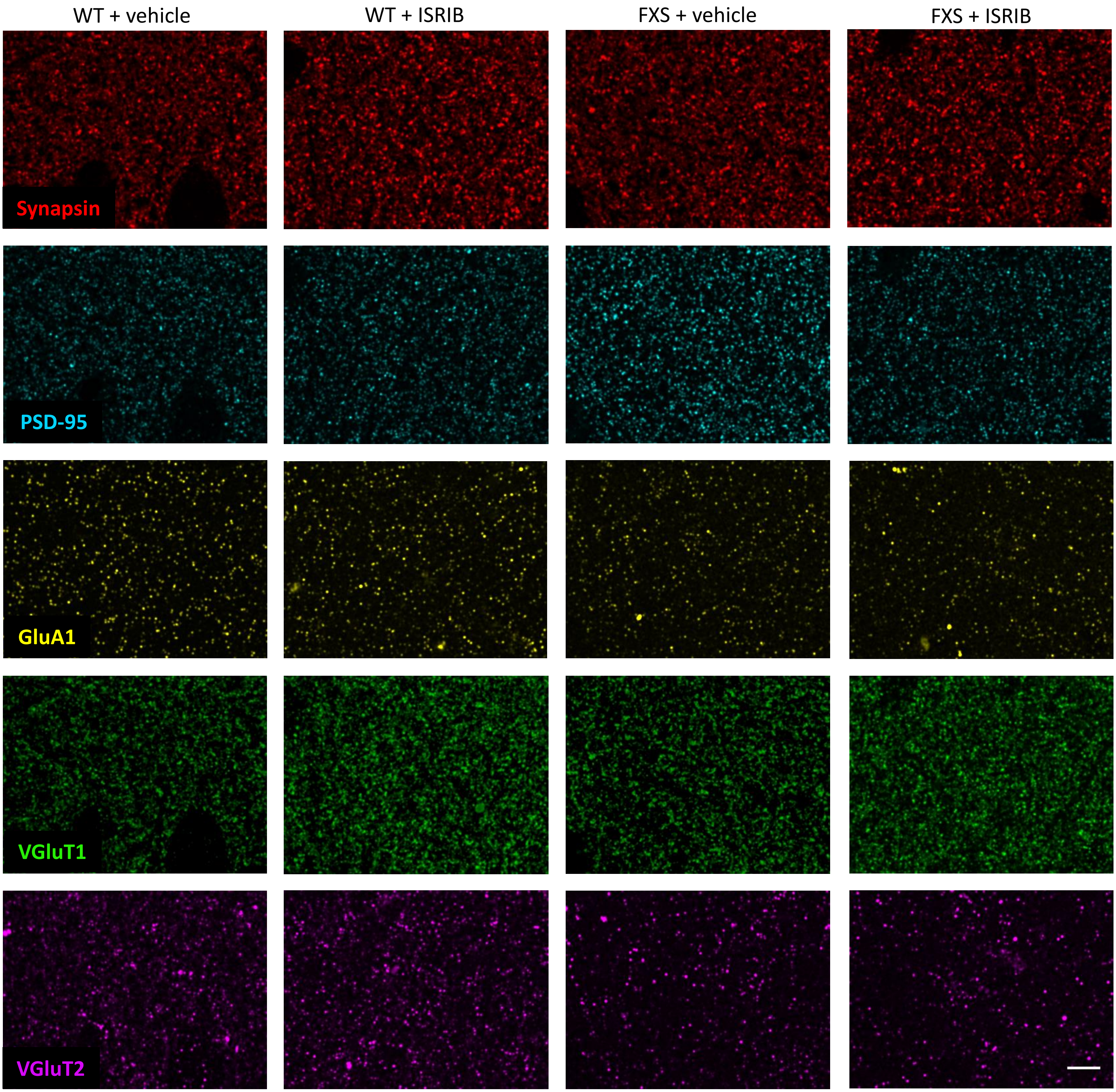

Supplementary Figure 3

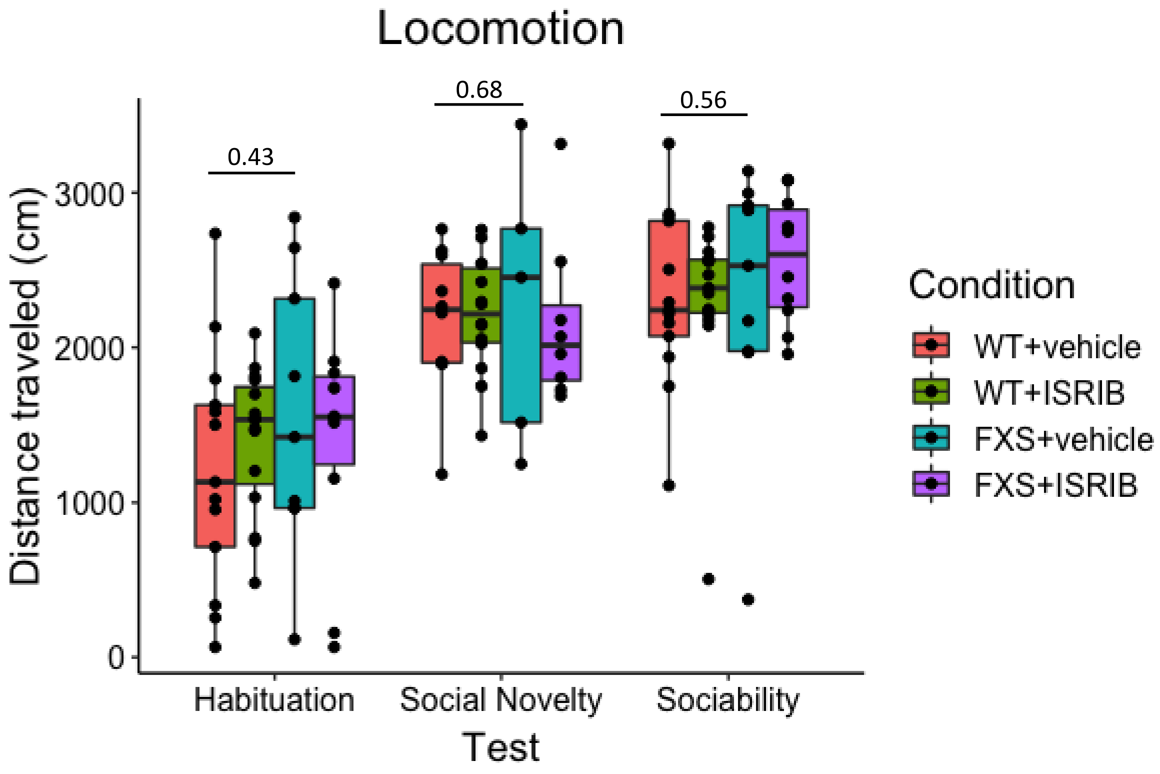

Supplementary Figure 4

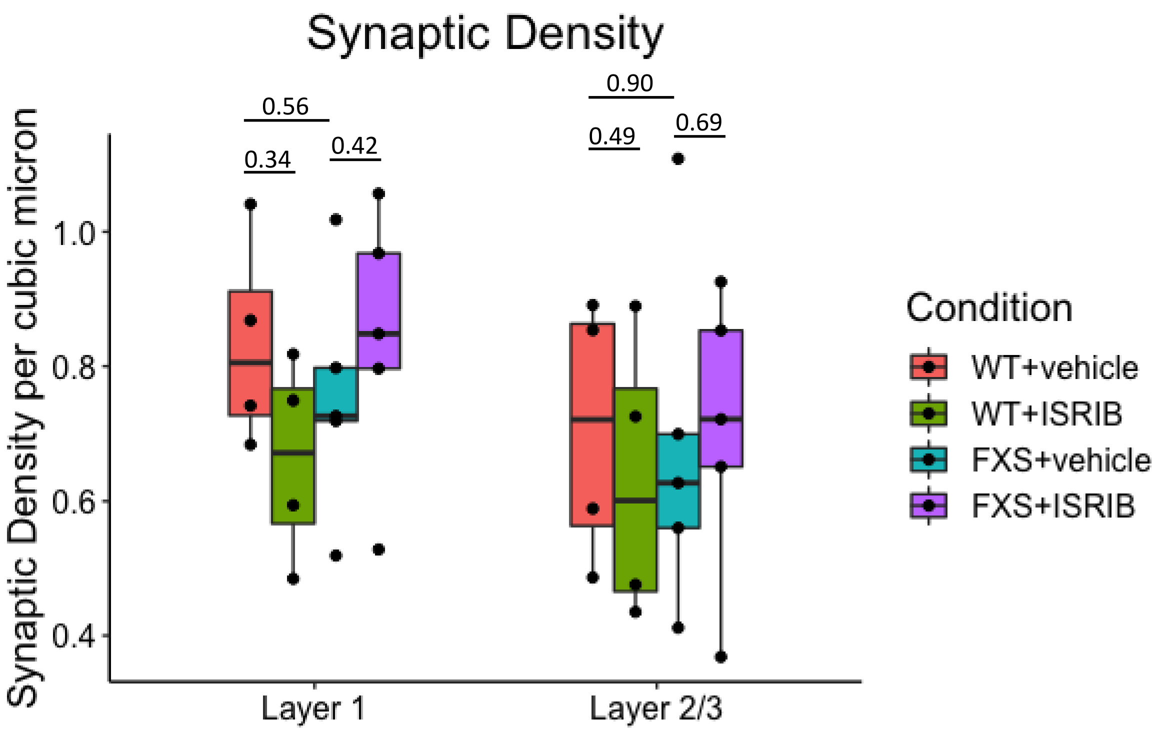
